# Distinguishing integrated sub-viral fragments from infectious Rubus yellow net virus in Scottish commercial red raspberry cultivars

**DOI:** 10.1101/2023.03.07.531471

**Authors:** Wendy McGavin, Sue Jones, Stuart MacFarlane

## Abstract

Short read sequencing of two field-grown plants of red raspberry cv Glen Dee identified sequences derived from Rubus yellow net virus (RYNV). Surprisingly, PCR primers designed to target these sequences amplified RYNV-specific DNA fragments from some high health nuclear stock raspberry plants, previously tested as free from RYNV infection. The complete sequence of a Scottish isolate of RYNV (RYNV LG) was determined and a panel of primers covering the entire virus genome was designed to demonstrate that nuclear stock plants of cultivars Glen Dee, Glen Ericht, Glen Fyne and Glen Moy all contain partial fragments of the RYNV genome as integrated elements but do not contain the entire RYNV genome.

The complete, circular genome of RYNV could be detected in RYNV LG-infected plants by rolling circle amplification but was not detected in Glen Dee plants. Grafting experiments showed that infectious RYNV could be transferred from RYNV LG plants but not from Glen Dee or Glen Moy plants, further demonstrating that these cultivars contain only partial, integrated RYNV elements. Grafting of RYNV LG to the certification-approved indexing species *R. occidentalis* (black raspberry) and *R. macraei*, showed that RYNV LG induced recognisable symptoms in both plants but that *R. macraei* was more susceptible and produced more recognizable symptoms. Effective testing for RYNV in high health certified raspberry plants can be achieved by combining graft testing with multifragment PCR.

## Introduction

Rubus yellow net virus (RYNV) infects wild and cultivated species of *Rubus* (*R. idaeus* [red raspberry], *R. occidentalis* [black raspberry] and *R. procerus* [Himalaya blackberry]) (Stace-Smith and Jones. 1987). Although there is very little information available on the economic impact of RYNV by itself on raspberry production (Stace-Smith, 1955), it is recognized that RYNV can be a component of the commonly occurring and widespread raspberry mosaic disease (RMD) complex that also includes black raspberry necrosis virus (BRNV) and raspberry leaf mottle virus (RLMV) (Converse *et al.*, 1987). These viruses are all transmitted by the same vector, the large raspberry aphid, *Amphorophora agathonica* (in the US) and *A. idaei* (in Europe).

Earlier work at the Scottish Crop Research Institute, Dundee (the predecessor of the James Hutton Institute) used heat treatment of a field-grown RMD-symptomatic red raspberry plant (cv Malling Jewel) to remove BRNV and RLMV and, thus, obtain an isolate of RYNV (Jones and Roberts, 1976). Examination of this plant by electron microscopy (EM) revealed bacilliform virus-like particles suggested to be RYNV, and this initial plant, and others grafted from it, are the sources of RYNV used in all subsequent RYNV studies at the James Hutton Institute. The bacilliform structure of RYNV suggested that it could be a badnavirus, which was later confirmed by amplification of a small portion of the RYNV genome using generic badnavirus primers, enabling the design of RYNV-specific PCR detection primers (Jones *et al.*, 2002). Badnaviruses are double-stranded DNA viruses with a genome size of 6.8-9.2 kb that, in common with other viruses in the *Caulimoviridae* family, replicate via reverse transcription of a terminally-redundant single-stranded mRNA (Bhat *et al.*, 2016). During this process sections of the badnavirus genome, and occasionally the complete genome, can become integrated into the host plant chromosomal DNA. In the case of three badnaviruses, Banana streak OL virus, Banana streak GF virus and Banana streak IM virus, and the caulimoviruses Tobacco vein clearing virus and Petunia vein clearing virus, activation of the integrated full-length genome can give rise to an episomal infection of the host plant (Bhat *et al.*, 2016). The presence of integrated badnavirus sequences in the host plant chromosomal DNA causes a problem when PCR amplification is used to assess if a plant is or is not infected with that virus.

To safeguard the quality and productivity of raspberry and blackberry crops, the UK and EU operate a certification scheme whereby mother plants that are used for the propagation of planting stocks are tested to ensure they are free from infection by specific pathogens, including RYNV (https://www.gov.uk/guidance/fruit-propagation-certification-scheme). The James Hutton Institute maintains a collection of commercially important red raspberry cultivars that are pathogen tested annually to comply with the high health certification requirements. The recommended testing method for RYNV high health certification is graft transmission to *R. occidentalis*, producing a net-like chlorosis of variable intensity, or to *R. macraei*, producing a much stronger vein netting and sometimes leaf distortion (Jones, 1991). Because *R. occidentalis* is also a recommended graft indicator plant for viruses other than RYNV, this species is the one most commonly used in certification testing.

Recent work has confirmed that isolated regions of the RYNV genome are integrated into the chromosome of several different cultivars of red raspberry (Diaz-Lara *et al.*, 2020), which makes the adoption of PCR for RYNV detection problematic. Because the original, published RYNV-detection primers were derived from only a small region of a single isolate, we sequenced a Scottish isolate of the virus and compared this with two other recently published RYNV sequences to design additional diagnostic primers. We have used these primers, in conjunction with biological tests, to investigate the RYNV-status of plants in the James Hutton Institute high health raspberry collection.

## Results

### Confirmation that RYNV isolate LG is a replicating, aphid-transmissible virus

A red raspberry plant of cultivar Lloyd George carrying RYNV is maintained in the James Hutton Institute soft fruit virus collection. This virus in this plant was transferred by grafting in the 1980s from the original RYNV-infected Malling Jewel plant described above and we have now designated this as the LG (Lloyd George) isolate of RYNV. As a first step in this work we confirmed that our stock collection Lloyd George plant contained replicating and infectious RYNV. We were able to use large raspberry aphids to transfer this RYNV isolate from Lloyd George plants to both Latham and Glen Prosen raspberry plants, where the transmitted virus was detected by PCR using the diagnostic RYNV primers (RYN1F and RYN1R) of Jones *et al.* (2002). No RYNV sequences were amplified from Glen Prosen and Latham control plants using these primers.

### Sequencing of the aphid-transmissible LG isolate of RYNV

At the start of the work only two apparently complete sequences of RYNV were published - Baumforth’s Seedling A (isolate BS; Accession No. NC_026238.1, Diaz-Lara *et al.*, 2015) and a Canadian (Ca) isolate (isolate Ca; Accession No. KF241951.1, Kalischuk *et al.*, 2013). These isolates have only 82 % nt sequence identity and isolate Ca was predicted to encode two open reading frames (ORFs) in one part of its sequence that were absent from isolate BS. We designed PCR primers to sequences from these RYNV isolates and used them and other primers designed during the sequencing work to amplify and clone thirteen overlapping fragments of RYNV LG covering the entire virus genome (Supplementary Fig. 1, Supplementary Table 1). Assembly of these sequences, obtained from at least three clones of each fragment, produced a circular molecule of 7837 bp (GenBank accession no. OK666525) which is 98.6 % identical to the RYNV BS isolate of 7836 bp. Similar to RYNV BS, the RYNV LG genome encodes on the plus strand ORF1 (210 amino acids; 23.9 kDa molecular weight, unknown function), ORF2 (153 amino acids; 16.9 kDa molecular weight, unknown function), ORF3 (1976 amino acids; 224.7 kDa molecular weight, contains domains for potential movement protein, coat protein, reverse transcriptase, RNAseH, protease), an ORF3-overlapping ORF4 (138 amino acids; 15.4 kDa molecular weight, unknown function) and on the minus strand ORF6 (69 amino acids; 7.2 kDa molecular weight, unknown function) (Fig. 1). A recent RYNV GenBank accession (MZ358192; RYNV-BiH) has a genome of 7816 bp and is also very similar (96.5 % identity) to RYNV LG, however, RYNV-BiH apparently does not encode an ORF1 due to a single base insertion causing a frameshift approximately 40 nt downstream of the potential ORF1 translation start site. The c. 7000 bp region of RYNV Ca, encoding ORFs 1, 2, 3, 4 and 6, is 81.5% identical to the same region of RYNV LG. Closer examination of the remaining RYNV Ca region, suggested previously to encode an ORF5 and an ORF7, show this to contain a 370 bp inverted repeat of part of ORF3 (Fig. 1).

**Figure 1.**
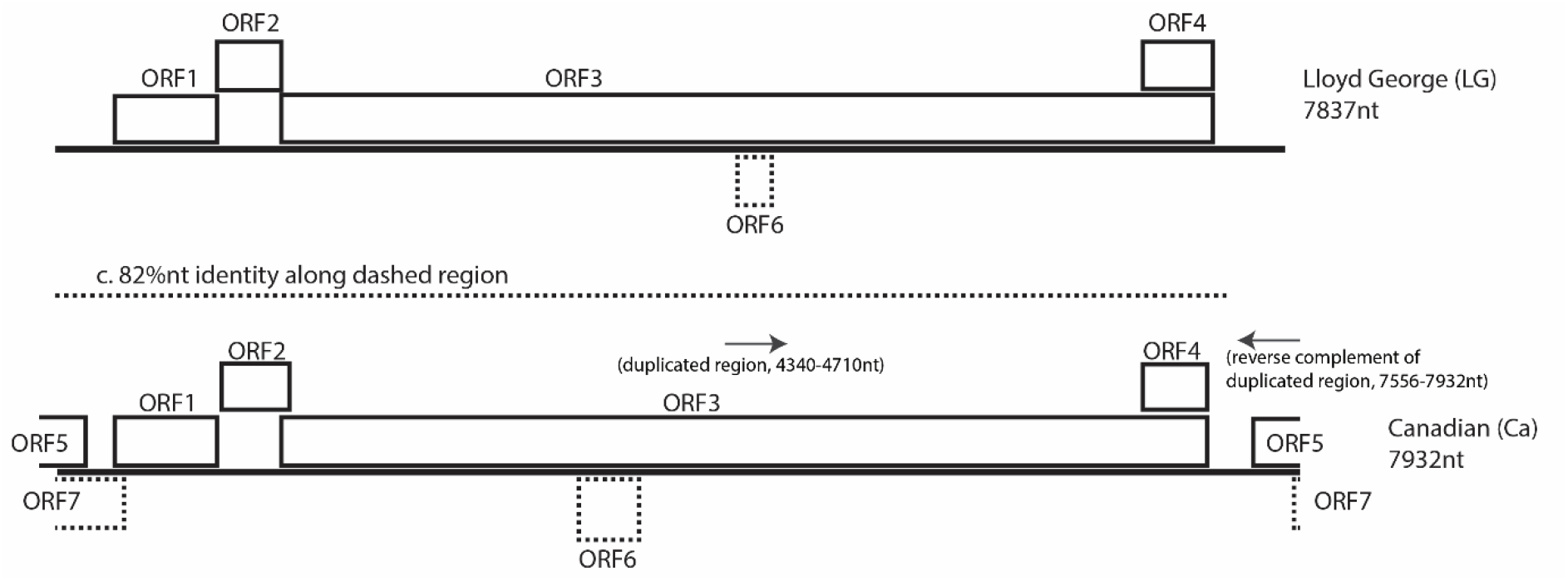
Comparison of genomes of RYNV LG and RYNV Ca. The circular DNA genomes are presented in linear form. Predicted open reading frames (ORFs) are shown as rectangular boxes. ORFs on the reverse DNA strand have dotted borders. The 3’ terminal sequence of RYNV Ca that is an inverse repeat of an internal ORF3 sequence and which gives rise to predicted ORFs 5 and 7 is denoted by an arrow.

### Identification of integrated RYNV sequences in Scottish commercial raspberry cultivars

Glen Dee is a commercially important raspberry cultivar produced at the James Hutton Institute and maintained as virus-tested mother plants in the JHI soft fruit high health facility. No symptoms of RYNV have ever been seen in these plants following grafting to the recommended indicator plant *R. occidentalis* and the plants test negative for RYNV using the original diagnostic RYNV primers (RYN1F and RYN1R) of Jones *et al.* (2002). In 2018 we did RNA short read paired-end sequencing of two field-grown Glen Dee raspberry plants showing strong yellowing symptoms and discovered sequences of several viruses including some apparently derived from RYNV (Supplementary Table 2). Two assembled contigs of 874nt from Glen Dee plant 5 (Dee_B5) and 2591nt from Glen Dee plant 6 (Dee_B6) overlapped one another, with the Dee_B6 contig also overlapping the c. 350nt region that would be amplified by the previously published RYN1F and RYN1R primers (Jones et al., 2002). Closer inspection showed that the Dee_B6 amplicon had six nucleotide differences to RYNV LG in these primer regions (Fig. 2) but that five of these six base differences were present in the RYNV Ca sequence of this region. Primers designed to the Dee_B6 sequence (3470/3471; Supplementary Table 1) were able to amplify RYNV-specific sequences from DNA extracted from both the Dee_B5 and Dee_B6 plants, showing the NGS sequence assemblies to be correct, but did not amplify RYNV LG (data not shown).

**Figure 2.**
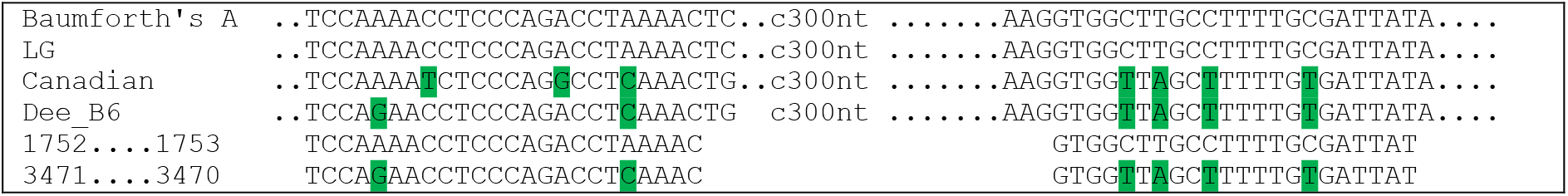
Nucleotide differences at diagnostic primer binding sites in RYNV sequences from the Dee_B6 deep sequencing data contig, published virus isolate genomes and PCR detection primers 1752, 1753, 3470 and 3471. Nucleotide differences are highlighted.

We initially thought that the Glen Dee and Lloyd George plants might be infected with different RYNV isolates that were sufficiently divergent to prevent them each being amplified using primers designed to the other isolate. To create a more “universal” RYNV PCR test we compared the two published sequences of RYNV available at that time (isolates BS (Genbank accession NC026238) and Ca (Genbank accession KF241951)) and designed further PCR primer pairs (3476/3477; 3478/3479; 3480/3481; 3482/3483) that were predicted to amplify fragments from both isolates. All four primer pairs produced amplicons of the expected size from both RYNV LG-infected plants and uninoculated Glen Dee plants from the pathogen-tested high health raspberry collection (Supplementary Fig. 2). Subsequently, one primer pair (3478/3479) was used to examine our entire collection of high health raspberry plants (that had previously tested as RYNV-free by grafting to *R. occidentalis* indicator plants). Although the majority of high health accessions tested negative for RYNV using these primers, amplified bands were obtained from the established Scottish cultivars Glen Dee, Glen Ericht, Glen Fyne and Glen Moy. The Glen Dee 3478/3479 amplicon (246bp) was cloned and sequenced, revealing it to have 88% identity to the RYNV BS isolate and 99% identity to the RYNV Ca isolate (Supplementary Table 2). However, this amplicon, which was located to the RYNV ORF1, had a single nucleotide deletion compared to the published BS and Ca sequences, which would cause a frame-shift mutation in the ORF1. This suggested that the Glen Dee amplicon was derived from a non-functional, integrated RYNV fragment very similar in sequence to the Ca isolate rather than from infectious RYNV in the plant. This result also suggested that Glen Ericht, Glen Fyne and Glen Moy might contain integrated RYNV fragments.

We then designed additional primers specifically to the RYNV Ca sequence between nucleotides 1224 and 3016 and used them to test DNA from high-health Glen Dee raspberry plants. The primer pair 4111/4106 amplified a 1252 nt fragment instead of the expected 1801 nt fragment (Supplementary Table 2). However, primer pair 4111/4110 (where primer 4110 is located between the 4111 and 4106 sequences in the wild-type RYNV genome) amplified a larger, c. 2.5 kb fragment instead of the expected 1376 nt fragment. Sequence analysis revealed that the 4111/4110 RYNV amplicon had a 549 nt deletion. The 4111/4106 amplicon revealed a duplication of the RYNV genome from nucleotides 1320 to 1668, and an additional duplication of RYNV sequences (genome position 2243 to 3024) inserted between the 1320-1668 duplication. These experiments showed that Glen Dee raspberry has at least two, different RYNV-derived inserted elements in its genome.

### Conserved pattern of RYNV inserts in Scottish commercial raspberry cultivars

Our studies described above showed that it was possible to design primers that differentiate between RYNV Ca (present in Glen Dee as integrated, recombinant sequences) and the functional, infectious genome of RYNV LG. We expanded this work to produce a panel of seventeen primer pairs that were expected to amplify either (a) RYNV Ca sequences alone, (b) RYNV LG sequences alone (c) both Ca and LG sequences together. These primers were used to test our high health cultivars Glen Dee, Glen Fyne, Glen Moy and Glen Ericht, and also RYNV LG in the original Lloyd George plant and when this virus was transferred by grafting into Glen Dee plants (Table 1).

**Table 1.** RYNV isolate-specific amplicon PCR from different raspberry cultivars. Amplicons detailed in unshaded rows are from infectious RYNV LG only. Amplicons detailed in light shaded rows are from both infectious RYNV LG and integrated elements. Amplicons detailed in darker shaded rows are from integrated elements only.

| Amplicon | Primer pair | Position of amplicon in RYNV genome | PCR amplification result |  |  |  |  |  |  |
| --- | --- | --- | --- | --- | --- | --- | --- | --- | --- |
|  |  |  | RYNV LG virus stock collection plant | Glen Dee high health plant | Glen Dee grafted with RYNV LG | Glen Ericht high health plant | Glen Fyne high health plant | Glen Moy high health plant | R. occidentalis cv. Cumberland |
| A | 4080/4081 | 7778 - 319 | + | - | + | - | - | - | - |
| B | 4078/4079 | 319-411 | + | - | + | - | - | - | - |
| C | 4082/4083 | 411 - 801 | + | - | + | - | - | - | - (ladder) |
| D | 3478/3479 | 788 - 1034 | + | + | + | + | + | + | nt |
| E | 4072/4073 | 1183 - 1443 | + | - | + | - | - | - | - (ladder) |
| F | 4109/4110 | 2184-2600 | - | + | + | + | + | + | nt |
| G | 4057/4058 | 2517-2834 | + | - | + | - | - | - | - |
| H | 4387/4389 | 2585 - 3090 | - | + | + | + | + | + | nt |
| I | 4387/4388 | 2585 - 3118 | - | + | + | + | + | + | nt |
| J | 4307/4308 epiBS-2604F epiBS-2715R | 2640-2715 | + | - | + | - | - | - | - |
|  |  |  |  | Glen Prosen | Autumn Bliss |  |  |  |  |
|  |  |  |  | - | - |  |  |  |  |
| K | 3482/3483 | 3878 - 4140 | + | + | + | + | + | + | - (ladder) |
| L | 4076/4077 | 4873 - 5339 | + | - | + | - | - | - | - |
| M | 4096/4097 | 4710 - 5474 | + | + | + | + | + | + | - |
| N | 4390/4392 | 5605 - 5895 | - | + | + | + | + | + | nt |
| O | 4391/4393 | 5876 - 6229 | - | + | + | + | + | + | nt |
| P | 1752/1753 RYN1F RYN1R | 6502 - 6848 | + | - | + | - | - | - | - |
|  |  |  |  | Glen Doll | Glen Prosen | Latham | Glen Ample |  |  |
|  |  |  |  | - | - | - | - |  |  |
| Q | 4107/4108 | 7034 - 7441 | + | + | + | + | + | + | nt |

nt; not tested. ladder; multiple bands of incorrect size (see Supplementary Fig. 3)

Eight primer pairs (amplicons A, B, C, E, G, J, L, P) amplified RYNV sequences only from the RYNV-infected Lloyd George stock plant or from Glen Dee plants that had been grafted with RYNV LG and, hence, were specific for infectious RYNV LG (Fig. 3). Five primer pairs (amplicons F, H, I, N, O) amplified RYNV sequences from Glen Dee, Glen Dee grafted with RYNV LG, Glen Ericht, Glen Fyne and Glen Moy but not from RYNV infected Lloyd George and, hence, were likely to be specific for one or more integrated RYNV elements that were shared between the four Scottish commercial cultivars. Four primer pairs (amplicons D, K, M, Q) amplified RYNV in all plants, identifying sequences that were shared between the integrated elements in the four tested commercial cultivars and RYNV LG.

**Figure 3.**
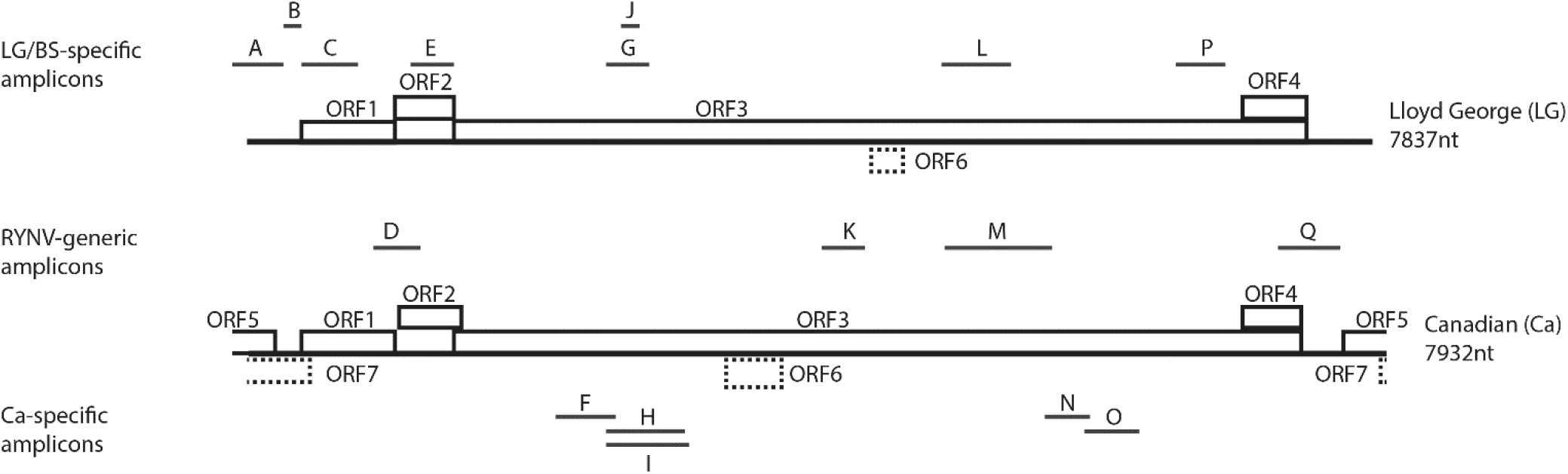
Distribution of detection primer amplicons across the RYNV genome. The upper row shows amplicons specific for RYNV isolate LG (demonstrated by this work) and RYNV isolate BS (predicted). The centre row shows amplicons generated from both RYNV LG and integrated sequence elements. The lower row shows amplicons generated from integrated sequence elements only (using primers designed to the Canada “isolate” sequence).

Interestingly, the original RYNV diagnostic primer pair (RYN1F/RYN1R; amplicon P) remained RYNV LG-specific and did not amplify sequences in Glen Dee, Glen Ericht, Glen Fyne, Glen Moy, Glen Prosen, Glen Ample, Glen Doll, Latham or black raspberry (*R. occidentalis*; Table 1., Supplementary Fig 3). Furthermore, primer pair 4307/4308 (epiBS-2604F/2715R; amplicon J), that was recently proposed as being specific for infectious (episomal) RYNV only (Ho *et al.*, 2021), in our studies also amplified only RYNV LG and did not amplify sequences in Autumn Bliss, Glen Dee, Glen Ericht, Glen Fyne, Glen Moy, Glen Prosen or black raspberry (*R. occidentalis*; Table 1., Supplementary Fig 3). These two primer pairs, used in combination, would be a very useful tool for initial screening of raspberry plants for potential infection by infectious RYNV.

### Rolling circle amplification of RYNV

Some badnaviruses have been detected by rolling circle amplification (RCA) which uses the bacteriophage phi29 DNA polymerase to amplify the circular badnavirus genomic DNA. Restriction digestion of the amplified DNA using enzymes known to recognise the genome sequence resolves the amplicons into fragments of predictable size. We detected the c. 8 kb circular dsDNA genome of RYNV LG using RCA of DNA extracted from the RYNV LG-infected Lloyd George plant. Digestion of the amplified DNA with NdeI produced the expected single fragment of c. 8 kb, and digestion with EcoRI produced the expected four fragments of c. 4200 nt, 1400 nt, 1250 nt and 900 nt (Figure 4A), showing this technique to be suitable for RYNV-detection. RCA of Glen Dee plant DNA produced a ladder of amplified DNA that did not resolve into RYNV-diagnostic fragments (Figure 4B) further confirming that Glen Dee does not carry replicating RYNV but only fragmented, integrated RYNV sequences.

**Figure 4.**
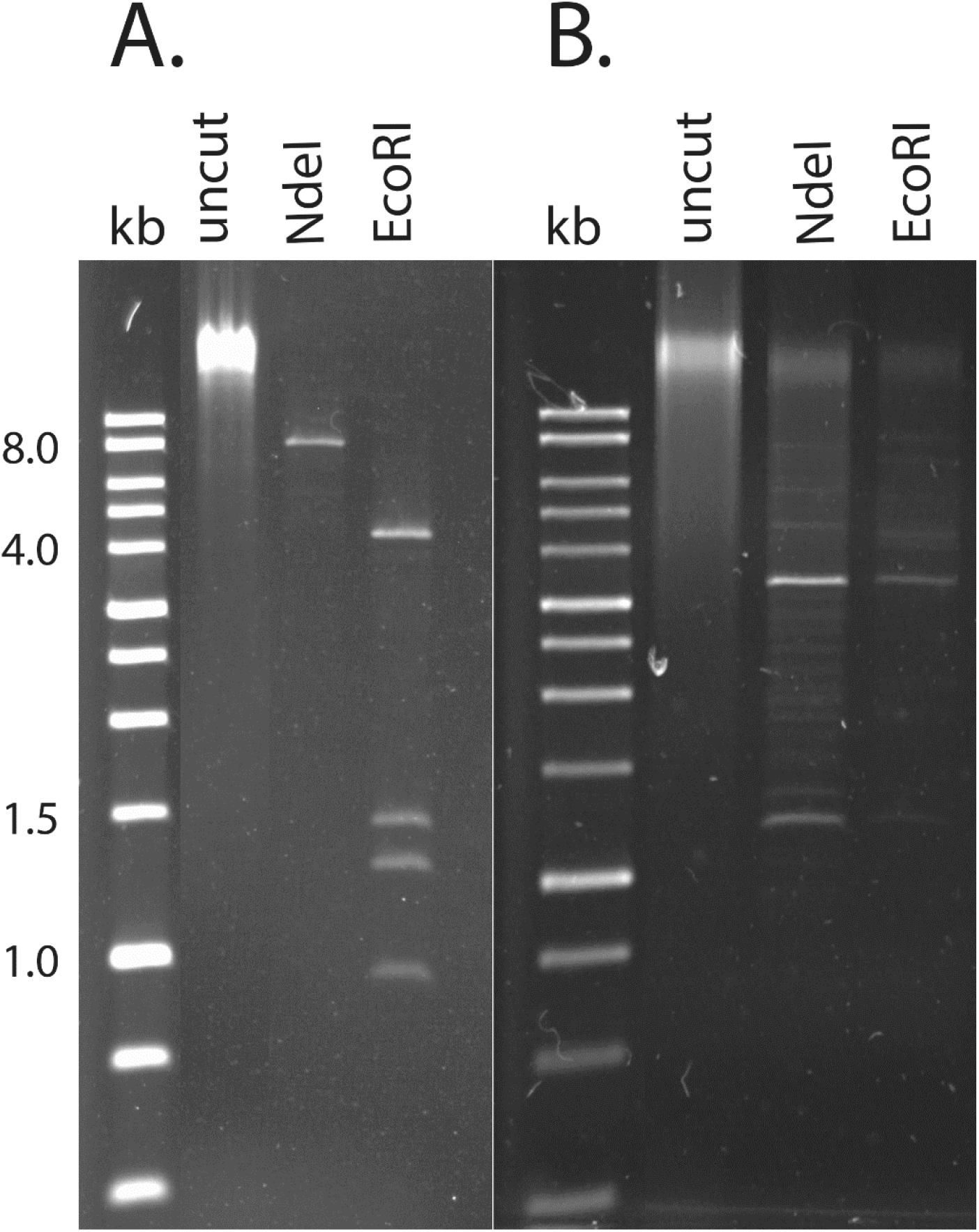
Rolling circle amplification of RYNV. Panel A. DNA from RYNV LG-infected Lloyd George plant. Panel B. DNA from a high health collection Glen Dee plant. Kb is 1kb DNA marker ladder (band sizes shown at left size). Uncut is RCA DNA with no restriction digestion. NdeI and EcoRI are RCA DNA digested with NdeI or EcoRI restriction enzymes.

### Symptom production after graft transmission of RYNV to *R. occidentalis* cv Cumberland and *R. macraei*

Because symptom clarity and reproducibility are essential elements in the use of grafting during high health plant testing, we examined the symptoms caused by RYNV LG in the two graft indicator species (*R. occidentalis* cv Cumberland and *R. macraei*). We also grafted Cumberland plants with scions from uninfected red raspberry cv Autumn Bliss plants to show that the grafting process itself does not induce any visible effects (Table 2). Four of five Cumberland plants grafted with RYNV LG tested positive for RYNV by PCR at eight weeks post grafting, however, only after 15 weeks did symptoms of chlorotic mottle develop in some leaves of these plants (Fig. 5). The fifth grafted plant remained uninfected by RYNV. For the three grafted *R. macraei* plants, extensive and intense areas of leaf blade chlorosis were visible on all plants after five weeks (Fig. 6) and all plants tested positive by PCR when assessed a week later.

**Table 2.** Graft transmission testing.

| Graft donor (scion) | Graft recipient (rootstock) | Number of RYNV PCR* positive plants/total grafted plants | Symptom development in graft recipient? |
| --- | --- | --- | --- |
| Lloyd George carrying RYNV LG | *R. occidentalis* cv Cumberland | 5/5 | 4/5; leaf and vein chlorosis |
| Autumn Bliss (uninfected, high health collection) | *R. occidentalis* cv Cumberland | 0/3 | 0/3 |
| Glen Dee (high health collection) | *R. occidentalis* cv Cumberland | 0/5 | 0/5 |
| Glen Ericht (high health collection) | *R. occidentalis* cv Cumberland | 0/3 | 0/3 |
| Glen Fyne (high health collection) | *R. occidentalis* cv Cumberland | 0/5 | 0/5 |
| Glen Moy (high health collection) | *R. occidentalis* cv Cumberland | 0/5 | 0/5 |
| Lloyd George carrying RYNV LG | *R. macraei* | 3/3 | 3/3; leaf and vein chlorosis |
| Glen Moy (high health collection) | *R. macraei* | 0/3 | 0/3 |

\*PCR tested using the episomal RYNV-specific primer pair 4307/4308 (epiBS-2604F/2715R)

**Figure 5.**
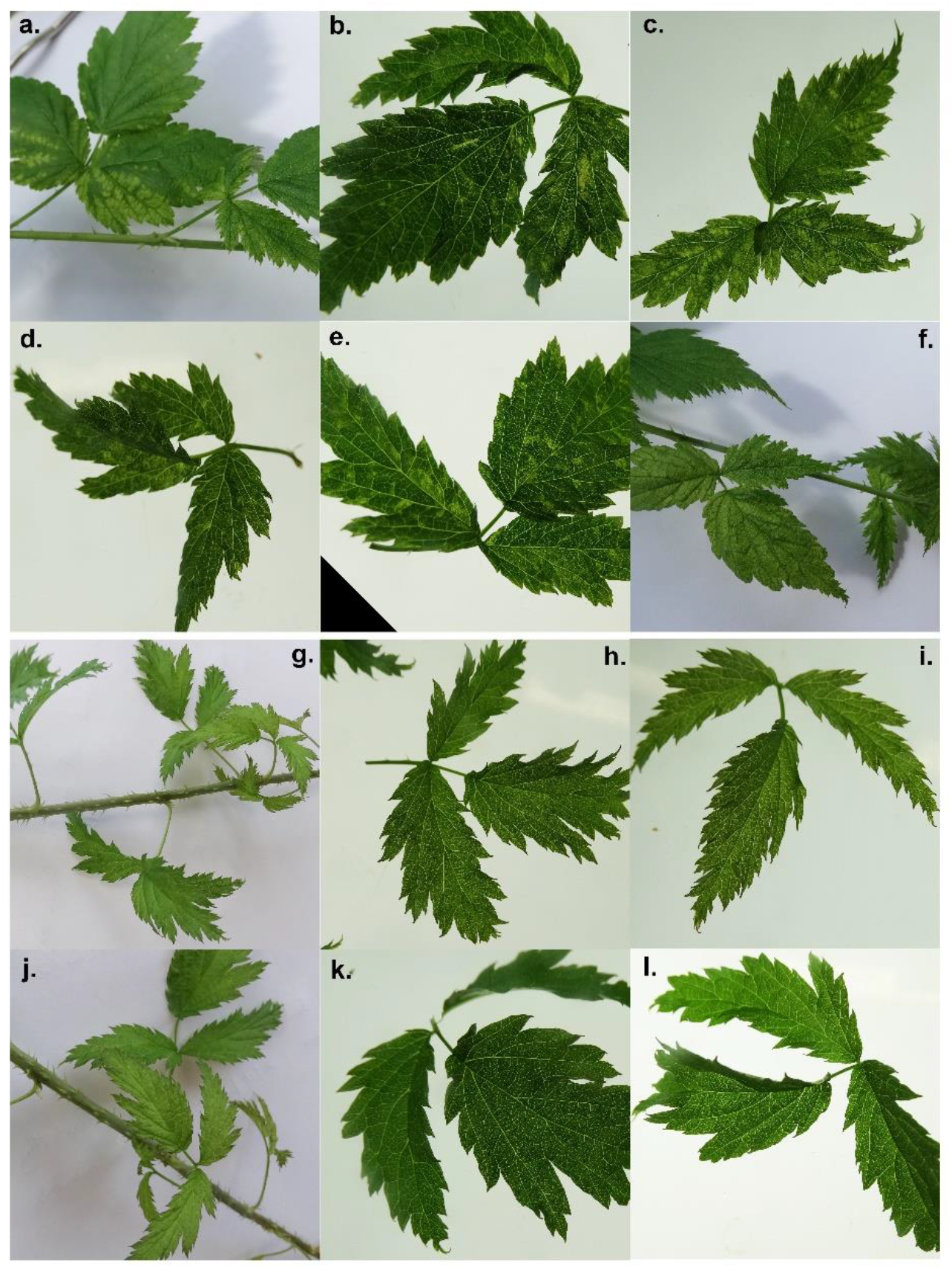
Symptoms on *R. occidentalis* cv Cumberland plants grafted with RYNV LG. Panels a. to f. RYNV LG. Panels G. to I. Cumberland plants grafted with healthy Autumn Bliss scions.

**Figure 6.**
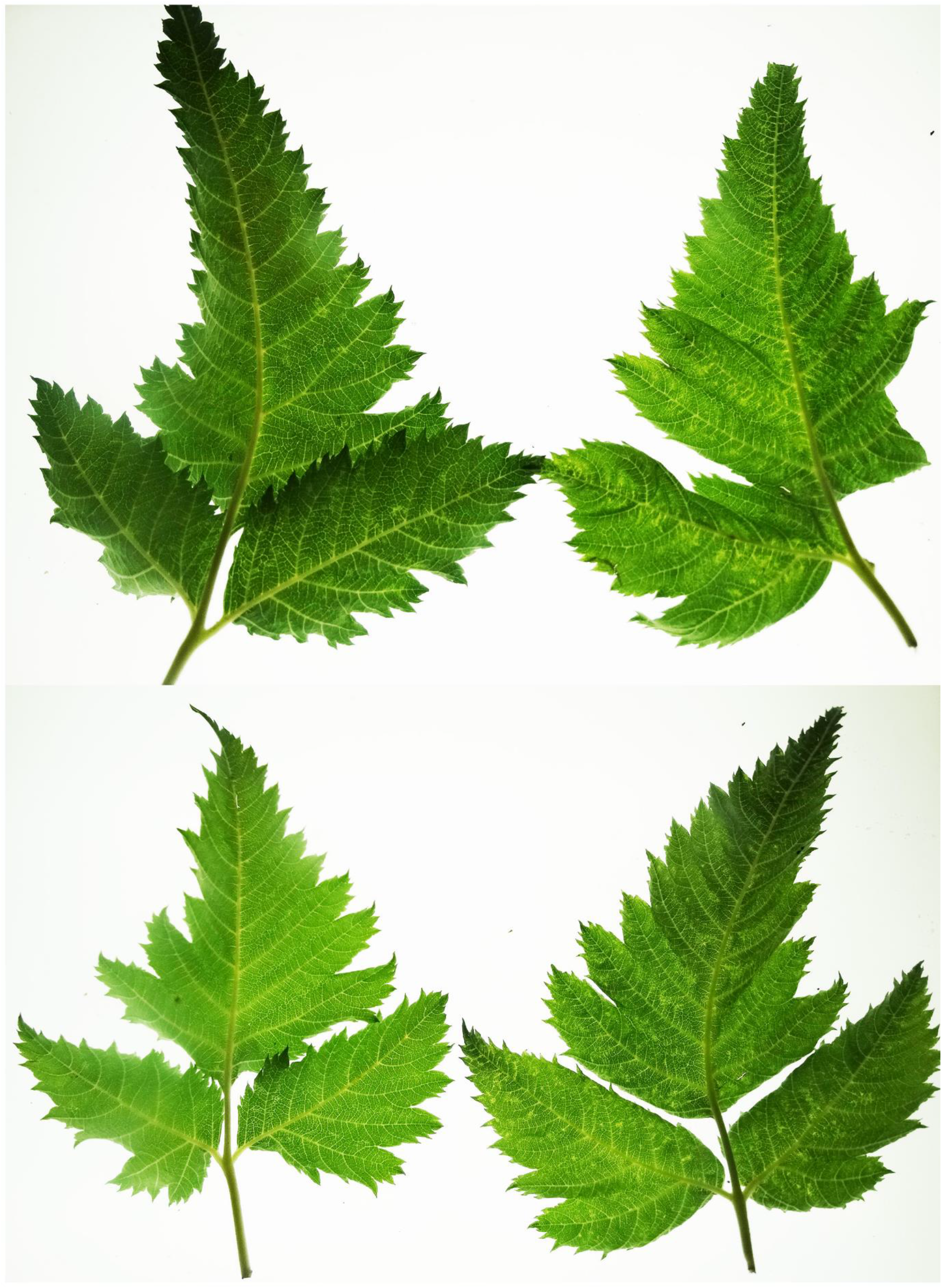
*R. macraei* plant symptoms after grafting with RYNV LG. Left panels are healthy (ungrafted) *R. macraei.* Right panels are RYNV LG-grafted *R. macraei*.

### Integrated elements in Glen Dee and other varieties do not release infectious RYNV and do not give resistance to RYNV

We grafted Cumberland plants with scions taken from Glen Dee (five plants), Glen Fyne (five plants), Glen Moy (five plants) and Glen Ericht (three plants). Over a four-month period no RYNV symptoms developed in any plant and PCR tests of all plants did not detect RYNV, showing that although all of the graft donors carried integrated RYNV sequences none of them could give rise to infectious RYNV.

To examine whether the integrated RYNV elements could give resistance to RYNV infection, perhaps by inducing post-transcriptional gene silencing, we challenged Glen Dee and Glen Moy plants by grafting with RYNV LG. PCR testing showed that all four Glen Dee plants became infected after graft inoculation. All PCR tests were negative at eight weeks post grafting, one plant was RYNV positive after 20 weeks and all four plants after 24 weeks. For Glen Moy, one of two plants became infected with RYNV by 24 weeks post grafting, although the virus was not detected at 20 weeks. These plants took longer to become detectably infected following graft inoculation than did the Cumberland plants that were grafted with RYNV LG, however, this could reflect a biological difference in susceptibility to RYNV between red raspberry and black raspberry rather than any protection derived from the integrated RYNV sequences present in the red raspberry plants.

## Discussion

In recent years much effort has been put into identifying, characterising and producing highly specific molecular tests for viruses that infect important fruit crop plants, including raspberry (Martin *et al.*, 2013). In most instances these tests are faster, cheaper and easier to carry out compared to the biological (grafting and mechanical transmission) tests that are long established but which take more time and require more extensive glasshouse facilities to complete. Although RYNV has been known about for nearly fifty years, a PCR test for the virus was only developed in 2002. Furthermore, an understanding that PCR amplification of RYNV sequences might be detecting integrated fragments of RYNV rather than infectious virus is very recent (Diaz-Lara *et al.*, 2020).

The original published RYNV PCR primers (RYN1F/RYN1R) were shown not to amplify any RYNV integrated sequences in uninfected *R. macraei* or red raspberry of the varieties Delight, Malling Jewel, Orion, Lloyd George, Glen Coe, Glen Clova, Glen Isla, Glen Prosen and Glen Rosa (Jones *et al.*,2002). We have now extended this set to include black raspberry (*R. occidentalis*) cv Cumberland and red raspberry Latham, Glen Ample, Glen Dee, Glen Doll, Glen Ericht, Glen Fyne and Glen Moy. Diaz-Lara *et al.* (2020) described a RYNV PCR primer pair (RYNV6-F/RYNV6-R) which did not produce an amplicon from black raspberry cv Munger but did amplify integrated elements in several red raspberry cultivars. Most recently, Ho *et al.* (2021) carried out a sequencing study of 33 historic red and black raspberry cultivars and examined them for the presence of integrated RYNV sequences. Six of the historic cultivars had no detectable RYNV sequences, whereas 27 cultivars did. In addition, no integrated RYNV sequences were found in black raspberry cv Munger. A pair of PCR primers (epiBS-2604F/2715R) was designed to the known infectious RYNV-BS isolate (Diaz-Lara *et al.*, 2015) targeting RYNV sequences that apparently are not included within any of the integrated sequences identified during the deep sequencing project. In our experiments, this primer set did amplify RYNV sequences from our RYNV LG stock collection plant and from Glen Dee when this plant had been grafted to contain RYNV-LG. Otherwise, the epiBS-2604F/2715R primer pair did not amplify any sequences from un-grafted Glen Dee, Glen Ericht, Glen Fyne, Glen Moy, Glen Prosen or *R. occidentalis* cv Cumberland. As the two primer sets, RYN1F/RYN1R (Jones *et al.*, 2002) and epiBS-2604F/2715R (Ho *et al.*, 2021), target different regions of the RYNV genome we propose to use both of these as an initial screen for RYNV testing in our certification procedure. We show here that RYNV can be detected by rolling circle amplification, however, it is an expensive and time-consuming technique requiring post-amplification restriction digestion. This may be appropriate for testing of new selections destined for registration as new cultivars but is less suitable for large-scale testing of plant collections.

The graft testing experiments showed that, as reported previously, *R. macraei* is the superior graft indicator plant for RYNV, producing strong visible symptoms very rapidly. *R. occidentalis* cv Cumberland does react to RYNV with symptom production, however, these symptoms are less clear than those of *R. macraei* and are much slower in their development. As *R. occidentalis* is also a recognized indicator plant for detecting other raspberry viruses it will probably continue to be the primary graft indicator species for raspberry certification testing. However, we suggest that PCR testing of plants using primer pairs RYN1F/RYN1R and epiBS-2604F/2715R, in combination with grafting to *R. occidentalis* would be a good approach to detect RYNV during certification testing.

## Acknowledgements

This work was funded by the Scottish Government’s Rural and Environment Science and Analytical Services division (RESAS).

## Methods

### Rubus grafting

Recipient plants were propagated in the JHI soft fruit high health facility from mother plants that were pathogen tested and free from RYNV. Donor plants were either RYNV-free from the high health facility or RYNV-infected and maintained in a separate licenced glasshouse. Scions from the donor plants were bottle-grafted with the graft junction sealed using parafilm. The plants were examined over a month in a heated (18°C) glasshouse to be certain that the grafts were successful, and after this were further maintained in the same glasshouse for virus infection to develop.

### Aphid transmission tests

A culture of large raspberry aphids (*Amphorophora idaei*) was established on red raspberry plants (cv. Glen Ample) derived from high health stock that was tested as being free from RYNV by grafting to *R. occidentalis* cv Cumberland. In addition, these plants did not produce a RYNV-specific amplicon when PCR tested with the RYNV-diagnostic primers RYN1F/RYN1R (JHI code numbers 1753/1752) (Jones *et al.*, 2002). Approximately 160 aphids were removed from these plants and starved for 60 minutes before being placed on the leaves of an RYNV LG-infected Lloyd George raspberry plant for 16 hours. Groups of twenty aphids were transferred to each of several RYNV-free Glen Prosen and Latham red raspberry plants and allowed to feed for 4 days at 18°C. The plants were treated with insecticide to kill the aphids and then transferred to a heated glasshouse. Leaf samples were collected 6 weeks later and tested by PCR for the presence of RYNV.

### PCR amplification of RYNV from raspberry

Frozen raspberry leaf samples were disrupted using a Qiagen TissueLyser and a stainless-steel ball bearing. DNA was extracted from the powdered leaf using the Machery-Nagel NucleoSpin RNA Plant kit according to the manufacturer’s instructions. PCR amplification of purified DNA was done using Phusion High-Fidelity DNA Polymerase (New England Biolabs). The integrity of the purified raspberry DNA was demonstrated by amplification of a 325nt portion of the EIF 2.1 gene using primers 1468(F) ATCATGAGGCCCTTCTGGAG and 1469(R) AACCATACCAGCATCTCCGT.

### Short read sequencing of Glen Dee raspberry

Glen Dee raspberry leaves (plants Dee_B5 and Dee_B6) showing vein yellowing were collected from a commercial raspberry plantation in Angus, north east Scotland. RNA was extracted from frozen, powdered leaf using the Qiagen RNeasy Plant Mini Kit using the provided RLT extraction buffer (450 μl) with the addition of Ambion Plant RNA Isolation Aid (45 μl) and β-mercaptoethanol (4.5 μl). The RNA was supplied to the Glasgow Polyomics facility for quality control assessment, ribosomal RNA depletion, library preparation and high-throughput sequencing using an Illumina NextSeq instrument (RD_PE2×75_33M). The raw RNA-sequence data for Dee_B5 and Dee_B6 has been deposited in the European Nucleotide Archive (ENA) and have samples accessions ERS2707341 and ERS2707342 respectively. The RYNV viral sequences in the two samples were identified using the bioinformatics workflow described in Jones et al (2019).

### Rolling Circle Amplification

Total DNA was isolated from RYNV-infected raspberry leaves using the Machery-Nagel NucleoSpin Plant II kit following the manufacturer’s instruction and extending the cell lysis and RNAse A step at 65°C to 1 hr to give a usual yield of c. 100-200 ng/μl. 1 μg of purified DNA was treated with 10 units Exonuclease V (New England Biolabs) in a 50 μl reaction at 37°C for 30 min to remove ssDNA, followed by heat inactivation at 70°C for 30 min. Rolling circle amplification (RCA) used the illustra TempliPhi 100 Amplification Kit. 5μl sample buffer was mixed with up to 5 μl nuclease-treated DNA and 1 μl of each of 4 RYNV-specific primers (3481, 3982, 3984, 3992; 20 μM [Supplementary Table 1]), which was heated to 95°C for 3 min to denature the dsDNA and then cooled in ice. 5 μl of reaction buffer and 0.2 μl enzyme mix was added and the reaction was completed at 36°C for 16 hr. Reaction products were examined with or without subsequent digestion using restriction enzymes predicted to cleave the RYNV genomic DNA.

## Figure Legends

Supplementary Figure 1. Diagram of overlapping PCR fragments used to clone and sequence the complete RYNV LG genome. The bold line is the complete genome in linear form, with left and right dotted lines denoting circularization of the genomic DNA. Numbers in red are forward primers and numbers in green are reverse primers. The thin lines represent the individual clones amplified by the flanking primers shown. Primers shown at the bottom of the figure were used for internal sequencing of cloned fragments.

Supplementary Figure 2. Amplification of RYNV-like sequences from RYNV LG-infected Lloyd George raspberry and uninfected (high health) Glen Dee raspberry plants using different primer pairs. Lanes 1-3; primers 3476/3477; lanes 4-6; primers 3478/3479, lanes 7-9; primers 3480/3481, lanes 10-12; primers 3482/3483. Lanes 1, 4, 7, 10 are no DNA; lanes 2, 5, 8, 11 are high health Glen Dee raspberry; lanes 3, 6, 9, 12 are RYNV LG infected raspberry. Kb is 1kb ladder marker (Promega).

Supplementary Figure 3. PCR testing of healthy R. occidentalis cv. Cumberland with RYNV-specific primers. RYNV isolate LG infected red raspberry tested with the same primers to show the amplicons produced from infectious virus. M1 is 1kb marker (Promega), M2 is 100bp marker (Promega). LG is RYNV isolate LG, C is Cumberland. Amplicon codes (A to P) are described in Table 1. Cumberland DNA was previously shown to be present and amplifiable using Rubus-specific EF1a primers
(Methods).

Supplementary Table 1. Primers used for complete RYNV LG genome cloning, sequencing and PCR detection.

Supplementary Table 2. Assembled contigs of RYNV-like sequences obtained by deep sequencing of field-grown Glen Dee raspberry plants and by PCR of a high health Glen Dee plant.

