## Supplemental Table 1 for "Distinguishing integrated sub-viral fragments from infectious Rubus yellow net virus in Scottish commercial red raspberry cultivars"

Supplementary Table 1. Primers used for complete RYNV LG genome cloning, sequencing and PCR detection

| Primer code | Sequence | Forward/reverse | Position in RYNV (5’ end of primer) |
| --- | --- | --- | --- |
| 1752 | TCCAAAACCTCCCAGACCTAAAAC | F (Jones et al., 2002) | 6502 |
| 1753 | ATAATCGCAAAAGGCAAGCCAC | R (Jones et al., 2002) | 6848 |
| 1766 | TGCTGACATCACACCAACAC | F | 7272 |
| 3470 | ATAATCACAAAAAGCTAACCAC | R | 6838 (Ca) |
| 3471 | TCCAGAACCTCCCAGACCTCAAAC | F | 6502 |
| 3476 | ATCACCGAGATCCACAATCAG | F | 644 |
| 3477 | AGCTGTTCGGCTACTTGTG | R | 939 |
| 3478 | GCCCTCAAGGCAGATCTG | F | 788 |
| 3479 | TCCTGCCACCTAGGGTTC | R | 1034 |
| 3480 | TCTACCTCAGGTACGAGGC | F | 2907 |
| 3481 | GTCTGCTGGTCCAATGGTG | R | 319 |
| 3482 | CTAAAAAGCCTGAGCTTCTGC | F | 3878 |
| 3483 | TCGTCTAGTCCCATGCCAT | R | 4140 |
| 3974 | ACCTACACCGGAGGACAGTTATC | F | 1126 |
| 4007 | CGTGTACTCCGAAGACCCGC | R | 3563 |
| 4010 | GACCTGCCTGTCATCGCTCC | R | 1949 |
| 4016 | TTCCTCGTAAGCGGCCATGAGCTT | R | 7338 |
| 4017 | GGGACTAGACGATAACTGGGATG | F | 4129 |
| 4018 | GTGGTGACTTTCTTGTAGATAGT | R | 5379 |
| 4019 | GTAGCCGAACAGCTGAGGAAGC | F | 926 |
| 4020 | CGACATCATGGCGTTGGGTAT | R | 2020 |
| 4031 | TCGATATGACACGAGGAGCGC | F | 1965 |
| 4032 | GACCTGCAACTCCTGATCCTC | R | 2959 |
| 4033 | CATGAAGGGAGGCGTAAGACT | F | 5320 |
| 4034 | TGCCTCTTCCGTGGGGAGTTT | R | 6541 |
| 4056 | TCTGAGGAGAGTACGGTGGA | R | 589 |
| 4057 | TCCGATATGACGCACCAGAA | F | 2517 |
| 4058 | GCTCCCTCTTTTCCTGGTCT | R | 2834 |
| 4062 | CGCTCTTGGTGTCTTTTGCT | R | 4568 |
| 4064 | TTCTTCGTCCATGAGCACCT | R | 5965 |
| 4065 | GATCTTCAACAGCCTTGCCC | F | 7173 |
| 4067 | AAGATGGACCAATGCTTCGC | F | 6062 |
| 4070 | GCTTTCACCATGGTAGCTTAAAC | F | 17 |
| 4071 | CGGAGATTTCGTGTGGTTGC | R | 407 |
| 4072 | GAATTATTGGTGCAAGTGCTAG | F | 1183 |
| 4073 | GGCTTCTGCTCATTGTATTGAG | R | 1443 |
| 4074 | CTGCAGGACAAGGCCGACTA | F | 3398 |
| 4076 | CGTAATCAAGGAAACACCAGAAA | F | 4873 |
| 4077 | GAGTCTTACGCCTCCCTTCA | R | 5339 |
| 4078 | TTGTAAGTGCGCCGATAGTGC | F | 7477 |
| 4079 | ATGCGGAAAAAGATGAACAAGTCT | R | 7758 |
| 4080 | TTCTTAAGTGTTCGAAAGCGCAC | F | 7778 |
| 4081 | CTAGTTGCTGTCGCTTTCTTAC | R | 319 |
| 4082 | TCGAAGCAGAGAGCAGCTCTT | F | 411 |
| 4083 | TCTGCCTTGAGGGCCTGTAG | R | 801 |
| 4092 | TCATCGGTAGTATGTCAGTCG | F | 1947 |
| 4093 | TGTCTTAGCCTGGCATCTGTG | R | 2319 |
| 4094 | GAAGGATATCTCACAAGTCCAG | F | 4357 |
| 4095 | CACGTCCTCCAGGTCATCC | R | 4732 |
| 4096 | TAGCGGATGACCTGGAGGA | F | 4710 |
| 4097 | AGTCATTGGCGTAGTACATGC | R | 5474 |
| 4100 | ACTACTCACCAATCCGAGCCA | F | 7091 |
| 4101 | TGAGCTTCCATGCTGCGGT | R | 7324 |
| 4105 | ACCACCAAGCTCTGCAAAAG | F | 2591 (Ca) |
| 4106 | CTTGCCCATAACTTCCTCG | R | 3022 (Ca) |
| 4107 | TGTTTTCTCGAGACTGCCCT | F | 7034 (Ca) |
| 4108 | GATGCTGCCGACACTAAAGG | R | 7441 (Ca) |
| 4109 | ACTACCTAGCAAGCAACGGT | F | 2203 (Ca) |
| 4110 | CTTTTGCAGAGCTTGGTGGT | R | 2601 (Ca) |
| 4111 | GCAGGCAATTGTCAACCTCA | F | 1224 (Ca) |
| 4112 | GGGTACTAGGTTCGCTCTCC | R | 1682 (Ca) |
| 4230 | GTCAAACTCCCAGAAAGTGTC | F | 1291 (Ca) |
| 4307 | TCCTACGAGGTAAGCCTAAGAG | epiBS-2640F (Ho et al., 2021) | 2640 |
| 4308 | CCGAGTTCGAGAGTTGGTTTAG | epiBS-2715R (Ho et al., 2021) | 2715 |
| 4387 | CCAAAACCACCAAGCTCTGC | F | 2586 (Ca) |
| 4388 | TGGGCTTGGTGGGATGAATC | R | 3134 (Ca) |
| 4389 | CCACCGGTAATGCTGCTGGA | R | 3106 (Ca) |
| 4390 | AATCAGCAGTAAACCCCCGG | F | 5627 (Ca) |
| 4391 | GCAATCGGCAATGCGAAGAT | F | 5898 (Ca) |
| 4392 | ATCTTCGCATTGCCGATTGC | R | 5917 (Ca) |
| 4393 | GTCGATCTCGGTTGCTGCTA | R | 6251 (Ca) |

Ca denotes RYNV Canada “isolate”-specific
