## Supplemental Table 2 for "Distinguishing integrated sub-viral fragments from infectious Rubus yellow net virus in Scottish commercial red raspberry cultivars"

Supplementary Table 2. Assembled contigs of RYNV-like sequences obtained by deep sequencing of field-grown Glen Dee raspberry plants and by PCR of a high health Glen Dee plant.

>Dee_B5_RYNV_length=874

ACAATCCAAACAACCCTGGAGCCCCAAAAGATATCTCTGCTCCGCGCAGAAGCAGAAGTC

GGAGAAGAGATCGAGCGTATGTACTACGCAAATGACTACTCTGAAGAAGGAGTCAGTCGC

CTGAGAAACCACAAACTGCTGCAGGAACTAAAAGAACAAGGCTACATAGGCGAAGAGCCA

ATGAAGCACTGGGCGAAAAACGGGATCAAGTGTAAGCTTGACATCAAGAACCCAGACATA

GTAATCAGCAGTAAACCCCCGGATGCTGTCTCAAAGGAGACGAAGGCACAATACCAGCGG

CACATTGACGCTCTCCTGAAGATCAAAGTAATCCAGCCAAGCAAGAGCAAGCACAGAACC

GCAGCCTTCATCACAAACTCGGGCACAACCGTTGACCCGATCACAAAGAAAGAAATCCGA

GGAAAAGAAAGGATGGTGTTCGACTACAGAAGTCTGAACGACAACACCCACAAAGACCAG

TATACTTTGCCTGGGATCAACACCATCATATCGGCAATCGGCAATGCGAAGATCTTCAGC

AAATTTGATCTGAAGTCTGGATTCCACCAAGTATTGATGGACGAAGAATCCATCCCGTGG

ACCGCATTTGTCACACCAGTAGGGTTCTACGAGTGGAAGGTAATGCCTTTCGGACTCGCA

AACGCTCCGGCCGTCTTCCAGAGAAAGATGGACCAGTGTTTTGCAGGAACCTCAGAGTTC

ATAGCCGTCTACATCGACGATATCCTGGTCTTCAGCAAGACCTTGAAGGAGCACGAAAAG

CACCTGAGCATCATGCTTGGGATATGTCGAGACAACGGCCTGGTTTTGTCACCAAGCAAG

ATGAAGTTAGCAGCAACCGAGATCGACTTCTTGG

>Dee_B6_RYNV_length=2591

GATGTATACGGCGTCGCTGATGTTTTGTGCAAGATCGTCGAAGACCTTGATTCTTCGCTG

CATGGCCACTCTTCGGGATTCGTCCATCTCCGCCATAGTACATGGTGCGGCTCTGATGGG

CGAGTAGTCACGGATTTCTCCGCTTGGAGGAGGGCAGTCTCGAGAAAACATGAGATGTTC

CTCTGTACCTCCTTGTTCACCAGGGCTTCGTCCCTTCCTTTGACCAGTTCGGCCGTGAGC

TCCGCTAGTTCCTTGGGAAGCCATCTTACTACAGTGATGAGTTGGGTAAGGCGACTAAGA

TTGTCAGCGAGCTGATTATCTTTGCCTTTGATGTGTTCGAACTTCATCTTCACCCCTGAG

TTTGTTATATAATCACAAAAAGCTAACCACCTTACCCGAGAAGGTTTCTTGGCGTTTAAC

TTTTCGTAAAAGGTGATGATGGCGTGGCAATCGGTTCTGATGGTGATCTCGTCCTTATTC

ATGTAGAAAATTTTGAATTTTTCTAAGGACTCCATTACCGCGAAGATTTCTGCGTCAATG

GTAGATTTCACTGTCGGGAACTTACCGCTTGCGTATGCGCAGATTTCTTCCTTGCCAGCT

GAGTCTGCTTTGTTGGGCTTCCACTTACAGACTCCGCCCCAACCTTCCATACAACCATCT

GTTTCGATGATCATATAGGCCTCCTCACTGGGCAGTTTGAGGTCTGGGAGGTTCTGGACC

AGGCTTTTGATCTTCTTTACTAAGGCCCAATCCGATGCATGCCATCTTCTGTCTCCATGC

TCGCTGGTCTTGCTGTATAGCGGGCCTAGGAGTGTTCCGCACTTCGGGATGTAGTTCCTG

GCATAGTTGAGAACTCCCAACCAACTTCTCAACCCTTTGAGGGTTTTTAGAGATTCATCG

TCCACCTCAGCTATCTTCTTGATTATGTGAGGCTGGAGTTTAATCTTTCCGTCACCAATG

GTGGCTCCCAAGAAGTCGATCTCGGTTGCTGCTAACTTCATCTTGCTTGGTGACAAAACC

AGGCCGTTGTCTCGACATATCCCAAGCATGATGCTCAGGTGCTTTTCGTGCTCCTTCAAG

GTCTTGCTGAAGACCAGGATATCGTCGATGTAGACGGCTATGAACTCTGAGGTTCCTGCA

AAACACTGGTCCATCTTTCTCTGGAAGACGGCCGGAGCGTTTGCGAGTCCGAAAGGCATT

ACCTTCCACTCGTAGAACCCTACTGGTGTGACAAATGCGGTCCACGGGATGGATTCTTCG

TCCATCAATACTTGGTGGAATCCAGACTTCAGATCAAATTTGCTGAAGATCTTCGCATTG

CCGATTGCCGATATGATGGTGTTGATCCCAGGCAAAGTATACTGGTCTTTGTGGGTGTTG

TCGTTCAGACTTCTGTAGTCGAACACCATCCTTTCTTTTCCTCGGATTTCTTTCTTTGTG

ATCGGGTCAACGGTTGTGCCCGAGTTTGTGATGAAGGCTGCGGTTCTGTGCTTGCTCTTG

CTTGGCTGGATTACTTTGATCTTCAGGAGAGCGTCAATGTGCCGCTGGTATTGTGCCTTC

GTCTCCTTTGAGACAGCATCCGGGGGTTTACTGCTGATTACTATGTCTGGGTTCTTGATG

TCAAGCTTACACTTGATCCCGTTTTTCGCCCAGTGCTTCATTGGCTCTTCGCCTATGTAG

CCTTGTTCTTTTAGTTCCTGCAGCAGTTTGTGGTTTCTCAGGCGACTGACTCCTTCTTCA

GAGTAGTCATTTGCGTAGTACATACGCTCGATCTCTTCTCCGACTTCTGCTTCTGCGCGG

AGCAGAGATATCTTTTGTGGTTCCAAGGTGGTCTGAATGGTGGTGACTTTCTTGTAGATC

GTCACTTCAGTACCTTCAAGCCTGACTCCACCTTTCATGGCTCGCAAGAAGTTGCAGCCG

ATGAGCATGTCTACGTCTGGTAGTTCCATTGGGAAGGCTGCAATGTACGGTGTATAGAAA

TCGCTTCCAGACACGATCATCTTACCGTTCTTCATTTTAAGCTTCGTTTCTCCAGTGGAA

TTTACTCCTGCGAAGTTCATCTGGTACTTCGCTTCCTCCAGGGCTTCTTTGGGGACTTTC

TTGCTGTTGATACATGTACAGGTGCATCCTGTGTCTATTACAGCTCTTACCCTGAACCTG

GTGGGTGCCCCCTTTTCCTGGGGTATTACAAACTCAACAATGGTGTTGTAGAGCTTGTTC

AATCCTCCTTTTGGCATTTCTGTGTATGCCGTGTCTACAGCTGCGGCTAGCTCCTCTATT

CCTTCTTCTGCTAAGAGGGCTCCAATAGTGTTGTTGCTACTTTCAGCTGTTTCTTTGATA

ATCACCTTGGGGGTGTCTTTCCACTGCTCGATGATCCTTGCCACGTTTTCCAAGTTTTGG

ATGTGGTCGATGGGAATCGTTGTTCCGAATTGGAAAGATTCCTTCTCCTTCCCCCTTTTT

GAGGCGTTGTCCAGGATGGACACGTCCTCCAAGTCATCCGCTAAGCTGATCAGGTCATCA

GGTTTGCGGTTGTGGAATTCCTGCAGCTGACTGGTAAGCTCCTCTACTTTCCTCGTGAGG

AAAGCGTTGTG

>Glen Dee_primer_pair_3478/3479 amplicon_length=246nt

GCCCTCAAGGCAGATCTGAAGGAAATCAAGGCACACAGTCAGCCCTGAAGCTAGGCTTCGCACAACTGCAGGAGGCAGTTCAGCTGATCATCACAAGGGAAAACGATCCCAAACCAATCGAAGCAGCTACTGCACAAGTAGCCGAACAGCTGAGGAAGCAGCTTATTGAGGTCAAGTCCGTCCTCGAGGAGACCAAGAAGATCGCGAGATCTCTGTCCCCCGACGGATGAACCCTAGGTGGCAGGA

>Glen Dee_ primer_pair_4111/4110 amplicon_c2500nt

GCAGGCAATTGTCAACCTCACAGCTCAGGTCACAAGGCTAGAAAAGACCGTCTCGGAGAAAGACACGGTCAAACTCCCAGAAAGTGTCCTCAACGACCTCACTAAGGAGTTCGGAAAGGTCAACTTAGGGAAAGGAAAGGGGATAGAAGGAGCAGTCTCATCCAGAGACAAGAACTTCTACGTCTGGAAGAACCCCTTCAATCAATACAATGAGCAGAAGCCAAACAAGGCTCCAGGCTCCGCCCGCAACTGAAAGGGCAACAAGCAGCTCGGAATCAGGCACCCCCACCTTGGAGGACCAGATCCGAGGATACAGTCGCTCCGCAAGGTTACGACACCAGGCGCAGCGAGCAATGAGAAGGACCTTCAGTAGGGACTTCAGAAACACCATAGAACGGCAACTAGACCCAGATGCCGAGCTTTCCCTCAGCAGAAGAAGGAGAGTAAACCGAGTACCAGCAGAGGTATACAGATCCGGAGAATGGGATTTACAACCCAGCAGGATCGTGGCACCACTAGCAGTCCCAACAGAAGCAAGGCTTAGCCAAAACAGGAATGGCAATATAAGCCTCAGATTCACCGACTTCCGAGATCAGAGGATCGTGGAGGAAGGAGAACCATCTGAGCCAGAAGGAAGgCCAGAAGGAGAAGATGATAGCACGCACTATGTGCTCATGTTCACCACTCAAGGTGGGACACCTTAGGGCACCAAGCGGGAAaTATNNNNNNNNNNNNNNNNNNNNN.....................................................................................................................................................................................................NNNNNNNNNNNNNNNNCTCaAGTTCAGACGAaGAaCCAAaGTTcGACGAGGCAGaGGACGaAGACGATGTgTACAACCAACAAaCCTgGCAAAaGGAaGACAaGGAGAAAAGAGACCTGGGACTACAGGGGTGGAAACCCACCGGGAGACCAGGAATCTACGAGATGATCCCCGAAGAAGAAGAAGAAATCCACTTCAGGTACGAGGCAGAGGACGAGGAGGATCAGGAAGTACAGGTGATAGGAGCTACCACCATTGAGGAGCCAGAGATGGAGTACCCAACTAGGCTCGAAGAAGTTATGGGCAAGCTCACTAAGGAGTTCGGAAAGGTCAACTTAGGGAAAGGAAAGGGGATAGAAGGAGCAGTCTCATCCAGAGACAAGAACTTCTACGTCTGGAAGAACCCCTTCAATCAATACAATGAGCAGAAGCCAAACAAGGCCCCAGGCTCCGCCCGCAACTGAAAGGGCAACAAGCAGCCCGGAATCAGGCACCCCCACCTTGGAGGACCAGATCCGAGGATACAGGCGCTCCGCAAGGTTACGACACCAGGCGCAGCGAGCAATGAGAAGGACCTTCAGTAGGGACTTCAGAAACACCATAGAACGGCAACTAGACCCAGATGCCGAGCTTTCCCTCAGCAGAAGGAGGAGAGCGAACCTAGTACCCGCGGAGGTACTATATGCACACAATGGCTCTGAGCCAGTAAACCGTGTGTACGAGCACTACAGTGAGCTCAGCGCTCATGTGGTAGATAGGCAGCAAGACTTCCGGTTCATCGAGGAAGTATCTTACCAGCACCTAGTCAGGGAAGGCATGCAATTTATACATGTCGGCATGGCGATGGTCAGAATCCAGATGCTGCACAGGACAGATGCAGGTATATCTGCACTAGTGGTGTTCAGAGACACCAGATGGAGTGATGACAGGCAGGTCATCGGAAGCATGTCCGTAGACATGACCAGGGGCGCGCAGTTGGTATACATAATATCAAACGCCATGATgTCAATACACGATTTCTACAATCGTATACAGGTCGGCGTGCAAATCCGAGGCTACGGAACAGGTTGGGAAGGTGGAGACAGTAACATGATCATCACGAGATCACTGGTTGGACGTCTCACCAACACCAGTATGACCAACTTCGAATACCGGATAGACCAGGTAACAGACTACCTAGCAAGCAACGGTGTGGCTTGCATCCCCGGTCAGAAGTGGAATGTGGCTAATAGATCTGGAGAATGGGAGTTACAACCCAGCAGAATCATAGCGCCACTAGTAGTCCCAACTGAAGCAAGGCTTAGCCAAAACAGAAATGGCAACATAAGCCTCAGATTCACCGACTTCCGAGATCAGAGGATCGTGGAAGAAGGAGGACCATCTGAGCCAGAGGGAAGGCCAGAAGGAGAAGACGATAGCACACACTATGTGCTTATGTTCAACCACTCAAGGTGGGACACCCTCGGCCAACCAAGCGGAAAATACGACTACATGGTACGCTACGATGCACCAGAACCAACCGCATGGCCCACAACCAACATCGGGTGGGACGATGATGAGTCACCAAAACCACCAAGCTCTGCAAAAG

>Glen Dee_ primer_pair_4111/4106 amplicon_length=1252nt

GGCAGGCAATTGTCAACCTCACAGCTCAGGTCACAAGGCTAGAAAAGACCGTCTCGGAGAAAGACACAGTCAAACTCCCAGAAAGTGTCCTCAACGACCTCACTAAGGAGTTCGGAAAGGTCAACTTAGGGAAAGGAAAGGGGATAGAAGGAGCAGTCTCATCCAGAGACAAGAACTTCTACGTCTGGAAGAACCCCTTCAATCAATACAATGAGCAGAAGCCAAACAAGGCTCCAGGCTCCGCCCGCAACTGAAAGGGCAACAAGCAGCTCGGAATCAGGCACCCCCACCTTGGAGGACCAGATCCGAGGATACAGGCGCTCCGCAAGGTTACGACACCAGGCGCAGCGAGCAATGAGAAGGACCTTCAGTAGGGACTTCAGAAACACCATAGAACGGCAACTAGACCCAGATGCCGAGCTTTCCCTCAGCAGAAGAAGGAGAGTAAACCGAGTACCAGCAGAGGTATACAGATCCGGAGAATGGGATTTACAACCCAGCAGGATCGTGGCACCACTAGCAGTCCCAACAAAAGCAAGGCTTAGCCAAAACAGGAATGGCAATATAAGCCTCAGATTCACCGACTTCCGAGATCAGAGGATCGTGGAGGAAGGAGAACCATCTGAGCCAGAAGGAAGgCCAGAAGGAGAAGATGATAGCACGCACTATGTGCTCATGTTCAaCCACTCAAGGTGGGACACCTTAGGGCAACCAAGCGGGAAATATGATTACATGGTGAGGTATGATGCACCAGAACCTACCGCATGGCCAACATCCAACATCGGATGGGATGATGATAAGCCACCCAAACCGCCAAGCCCTACAAAAGGATCTTTTGAGGTAAACCTCAAAGGAGAGAAGAAACTAAAAGAGAAGGAACTCGCGGAGTTCACACCGGAGACAGATCTGGTGAGCCAGTGGTTAAGTCAGCTGTCAACATCCGCACATAATAGCGGAGCCTCAAGTTCAGACGAAGAACCAAAGTTCGACGAGGCAGAGGACGAAGACGATGTGTACAACCAGCAAACCTGGCAAAAGGAAAACAAGGAGAAAAGAGACCTGGAACTACAGGGGTGGAAACCCACCGGGAGACCAGGAATCTACGAGATGATCCCCGAAGAAGAAGAAGAAATCTACCTCAGGTACGAGGCAGAGGACGAGGAGGATCAGGAAGTACAGGTGATAGGAGCTACCACCATTGAGGAGCCAGAGATGGAGTACCCAACTAGGCTCGAGGAAGTTATGGGCAAGC
