## Supplementary figures and images for "Distinguishing integrated sub-viral fragments from infectious Rubus yellow net virus in Scottish commercial red raspberry cultivars"

### Supplemental Figure 1

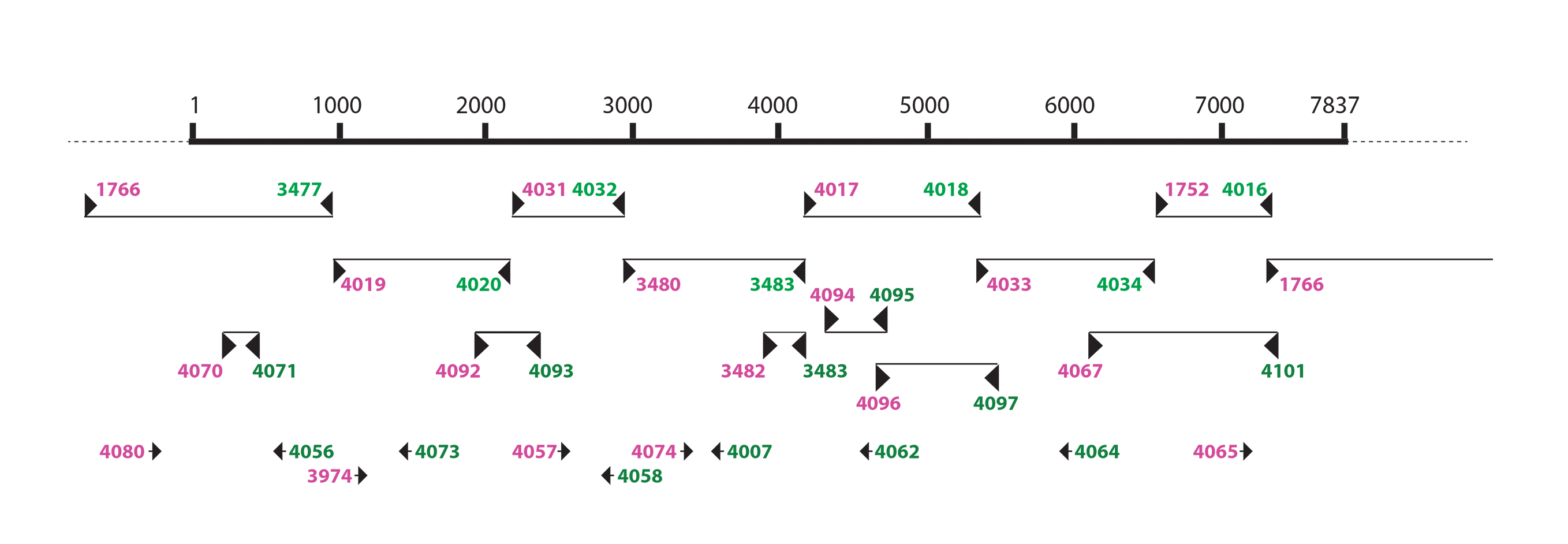

### Supplemental Figure 2

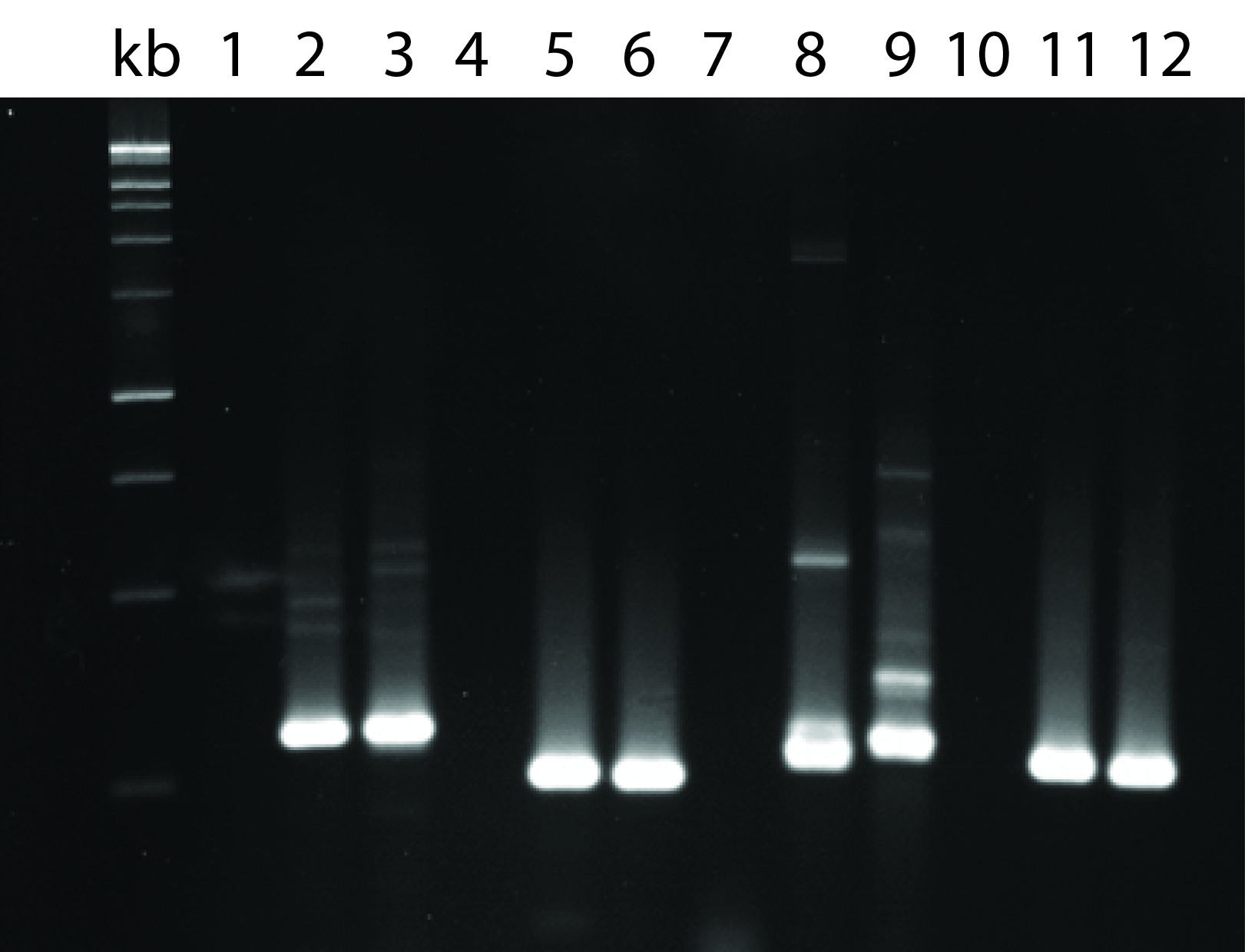

### Supplemental Figure 3

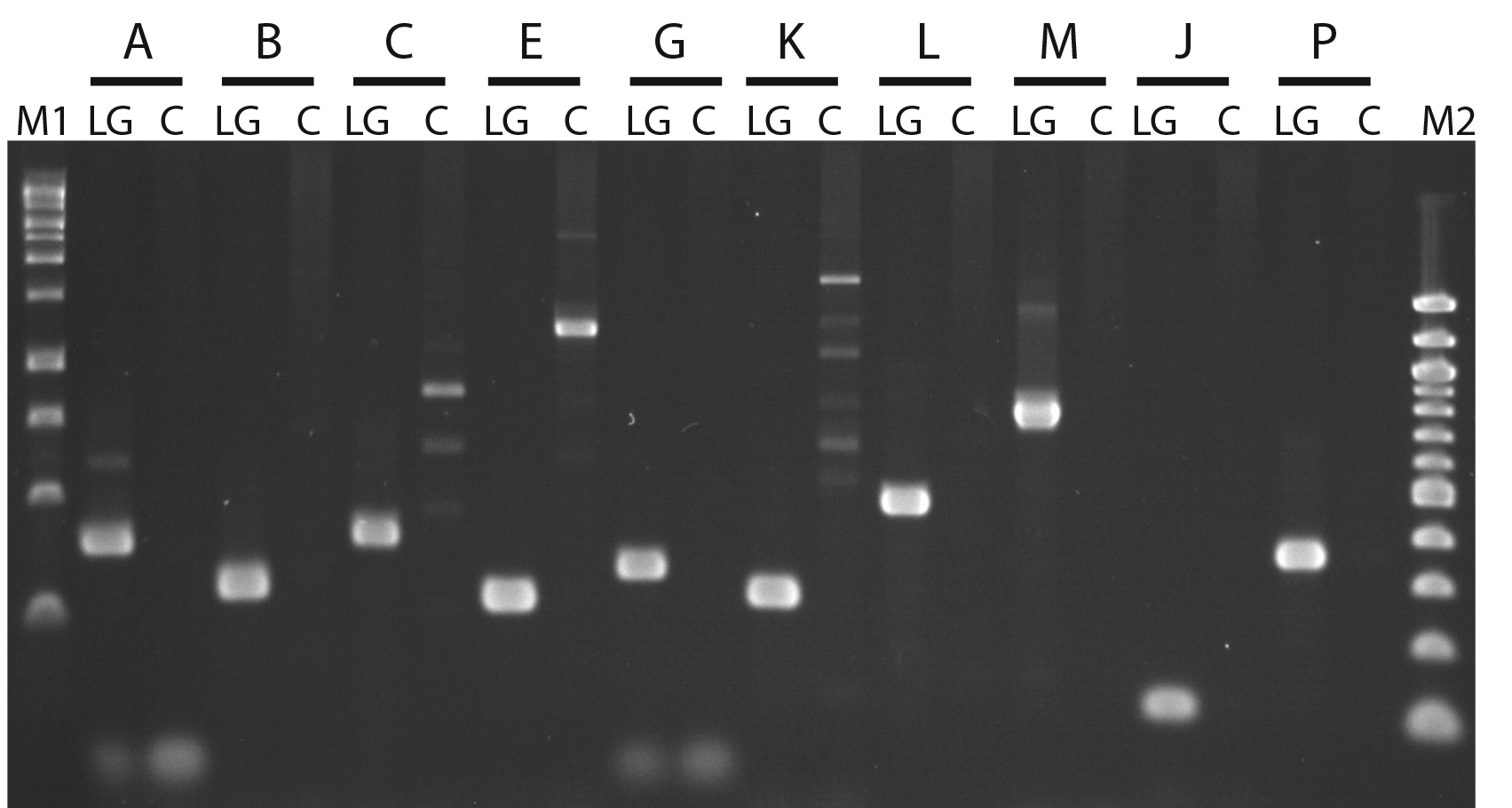
